# Hydrophobic bulk conservation and genetic code determine the pattern of prokaryotic genome organization

**DOI:** 10.1101/2024.01.09.574814

**Authors:** Wenkai Teng, Yangkai Zhou, Lingling Zheng, Yongqian Xu, Yangzhi Rao, Siruo Chen, Tian Xia, Wensheng Shu, Mengyun Chen

**Affiliations:** Shenzhen Key Laboratory of Marine Archaea Geo-Omics, Department of Ocean Science and Engineering, Southern University of Science and Technology, Shenzhen 518055, China; MOE Key Laboratory of Gene Function and Regulation, State Key Laboratory of Biocontrol, School of Life Sciences, Sun Yat-sen University, Guangzhou 510275, China; Department of Environmental Science and Engineering, University of Science and Technology of China, Hefei 230026, China; Guangzhou Key Laboratory of Subtropical Biodiversity and Biomonitoring, Guangdong Provincial Key Laboratory of Biotechnology for Plant Development, School of Life Sciences, South China Normal University, Guangzhou 510631, China

**Keywords:** hydrophobicity, genetic code, genome organization, natural selection, mutational biases, prokaryotes

## Abstract

Prokaryotic genomes in general exhibit marked organizational asymmetry, and differ substantially in GC-skew, AT-skew, and GS-bias (gene strand bias). Despite enigmatic origins, these organizational features and genomic GC-content have been described to be widely associated with each other recently, providing an opportunity to probe the evolutionary mechanisms. By analyzing sequence data of 4,012 leading strands from all available representative genomes, we found that the unusual nucleotide usage of coding genes under low GC-content contributed to a shift between two organizational patterns in the context of mutational biases. Analysis of artificially established neutral and natural models further suggests that in genes both GC-skew and AT-skew increase with decreasing GC-content, mostly contributed by synonymous substitutions because of genetic code, and by nonsynonymous substitutions because of hydrophobic bulk conservation in amino acid usage, respectively. This novel mechanistic framework in our study highlights the importance of evolutionary processes “operating at lower levels of organization” in the microbial world.

## Introduction

Prokaryotic genomes display intrinsic symmetries of organization in DNA sequences, e.g., Chargaff’s first parity rule, Chargaff’s second parity rule as well as theta-mode replication ^1,2^. However, there are also pervasive asymmetries in respect to nucleotide composition (usually measured by comparing the proportion of four nucleotide bases: adenine (A), guanine (G), cytosine (C), and thymine (T)) and coding gene distribution between the leading and lagging strands. Such asymmetries can be characterized by GC-skew (the proportional disparity between G and C in single-strand DNA which is calculated as (G - C) / (G + C) × 100), AT-skew (calculated as (A - T) / (A + T) × 100), and GS-bias (gene-strand distributional bias) ^3–5^. Although these organizational features of prokaryotic genomes have been described and investigated since 1996, the mechanisms account for them remain highly enigmatic ^6,7^.

The strand asymmetries in nucleotide usages were originally attributed to mutational biases caused by asymmetrical errors in DNA synthesis ^8^. Canonically, C and 5-meC can be spontaneously deaminated into uracil (U) and T with a higher possibility on the leading strand which serves as the template for lagging strand replication thus being in a single-strand state for a longer time ^9–11^. As a result, the leading strand has an excess of G over C (*i.e.*, GC-skew +) and an excess of T over A (*i.e.*, AT-skew -) in contrast with the lagging strand ^7,8^. The replication-driven cause of nucleotide skews has recently been studied experimentally in *Escherichia coli* using cytosine deaminase ^12,13^. According to such hypothesis, GC-skew should be negatively correlated with AT-skew and GC-content as the mutagenic deamination of C also leads to a net loss of G-C base pairs ^14^. Both GC-skew + and AT-skew +, however, have been observed in the leading strands of many genomes with low GC-content and particularly strong GS-bias (mainly belonging to the phylum Firmicute) ^15,16^. This atypical organizational pattern may originate from an unusual way of DNA replication with the participation of an additional polymerase named PolC ^17,18^. Unexpectedly, it has been demonstrated that non-synonymous sites of protein-coding genes in these genomes display stronger AT-skew, suggesting that the nucleotide usage of coding genes, likely shaped by some form of natural selection, as well as the gene distribution likely play an important role in determining the nucleotide skew pattern of prokaryotic genomes ^19^.

Indeed, coding genes are encoded preferentially on the leading strand (*i.e.*, GS-bias +) to avoid head-on collisions between the replication and transcription complexes which are very harmful to the replication fork ^20,21^. It has been reported that genes encoded on the lagging strands can accumulate more deleterious mutations ^22^. Nevertheless, such an interpretation seems mainly suitable to highly expressed genes and multi-gene operons, and the degree of GS-bias varies widely among different genomes ^5,23,24^. On the other hand, in some studies the nucleotide usage of coding genes was thought to be influenced by the nucleotide skew of strands due to separate codon and amino-acid usages of coding genes located on the leading and lagging strands ^25–27^. Regardless, strong GS-bias in many low-GC genomes possibly driven by horizontal gene transfer, and its significant correlations with GC-skew and AT-skew promoted researchers to propose that these organizational features should be mechanistically associated owing to a joint action of both mutational biases and natural selection ^16,28^.

In conventional ideas, adaptive processes operating at the population level account for everything in the microbial world ^29–31^. However, many ongoing efforts on mutational biases highlight the importance of nonadaptive processes operating at lower levels of organization than the population, particularly for the interpretation of phylogenetically conserved features which are tightly related with each other, like the above introduced GC-skew, AT-skew, GS-bias, and genomic GC-content ^32–35^. The correlations between them appear to show different organizational patterns, and several critical knowledge gaps remain regarding how organizational features are linked to genomic GC-content, the causal relationships between the distribution (and nucleotide usage) of genes and genomic nucleotide skews, and how natural selection shapes the nucleotide usage of genes. Here, to address these knowledge gaps we systematically explored all available complete sequences from prokaryotic representative genomes. In previous studies, the generally applied Z-curve method and recently developed tools like SkewIT mainly focus on evaluating the degree of skewness, but do not well quantify the degree of nucleotide disparities ^36–38^. Therefore, for better quantification of those organizational features we extracted the putative leading strands from complete sequences according to their Z-curves. GC-skew, AT-skew, GS-bias, as well as GC-content of these leading strands were then calculated and their associations were thoroughly investigated. Our results revealed two distinct organizational patterns based on the correlations of these features and contributed to a novel mechanistic framework for prokaryotic genome evolution, through which we confirmed decisive influences from hydrophobic bulk conservation and genetic code, which limits the amino-acid and codon usage at a low level of organization, on adaptative or nonadaptive processes.

## Results

### GC-skew, AT-skew, GS-bias, and GC-content are tightly associated

From 4,218 downloaded representative genomes, a total of 4,012 sequences of leading strands with positive values of GC-skew were obtained, including 3,481 chromosome and 531 plasmid sequences (Supplementary data 1 & 2, see Methods). Analysis of the sequences demonstrates that all correlations between GC-content, GC-skew, AT-skew, GS-bias, and sequence length are significant (Fig. 1A). In detail, GC-skew, AT-skew, and GS-bias are positively correlated with each other, with correlation coefficients ranging from 0.47 to 0.74, but all are negatively correlated with GC-content and sequence length, with correlation coefficients ranging from -0.21 to -0.71 (Fig. 1A). Of these, the strongest positive correlation is found between AT-skew and GS-bias, and the strongest negative correlation is observed between GC-skew and GC-content. To rigorously test these associations, sequences from bacteria and archaea, as well as chromosomes and plasmids, have been analyzed separately, all showing remarkably congruent patterns (Fig. S1). The results indicate a robust association among GC-skew, AT-skew, GS-bias, and GC-content of prokaryotic complete sequences.

**Figure 1.**
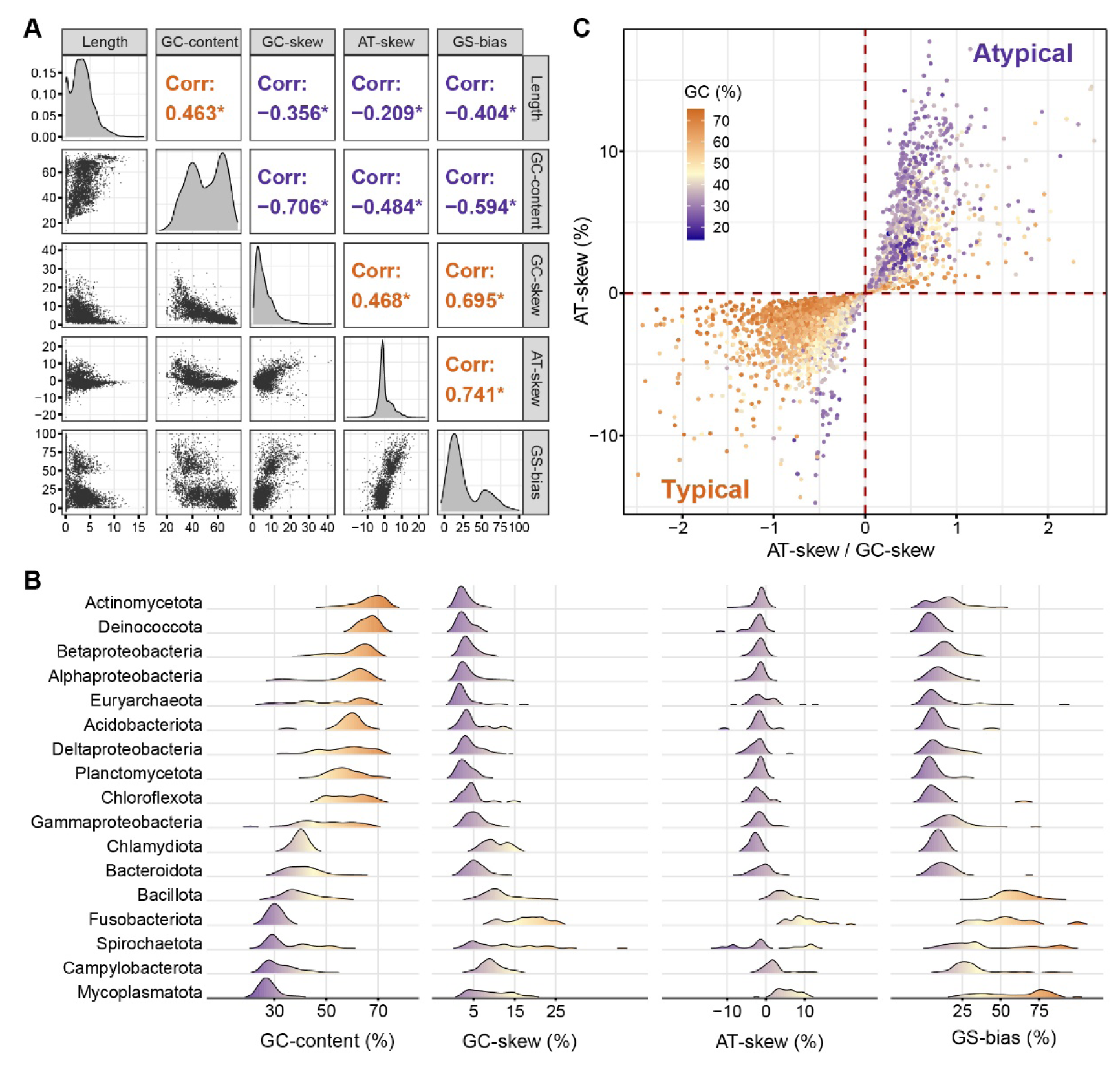
Relationships among different organizational features and GC-content of all leading strands. (A) Pairwise correlation coefficients between GC-content, GC-skew, AT-skew, GS-bias, and sequence length. Asterisks indicate adjusted p-value < 0.05. (B) Distributions of GC-content, GC-skew, AT-skew, and GS-bias in different phyla. (C) The values of AT-skew and AT-skew / GC-skew indicate two distinct organizational patterns.

Furthermore, distributions of these features across different lineages gave consistent results (Fig. 1B). High-GC lineages, mainly including Actinomycetota, Deinococcota, Pseudomonadota (formerly part of the phylum Proteobacteria, consist of Alpha-, Beta-, and Gamma-proteobacteria), Euryarchaeota, Acidobacteriota, Planctomycetota and Chloroflexota are characterized by relatively low GC-skew, AT-skew, and GS-bias. Whereas low-GC lineages, including Bacillota (formerly Firmicutes), Fusobacteriota, Campylobacterota, Mycoplasmatota and some lineages in Spirochaetota are mainly characterized by relatively high GC-skew, AT-skew, and GS-bias. GS-bias in these low-GC lineages can vary to more than 75% (Fig. 1B). These organizational features of prokaryotic genomes, as is the case for genomic GC-content, are pretty conserved likely resulting from early evolution ^33^.

### AT-skew and GC-skew display two distinct organizational patterns

As described in previous studies ^2,4,19^, typically the leading strand displays GC-skew + and AT-skew -, as is the case for *Pseudomonas furukawaii* (Fig. S2), in contrast with the atypical organization where the leading strand displays GC-skew + and AT-skew +, as is the case for *Shouchella miscanthi* (Fig. S3). Therefore, all complete sequences can be clearly classified into two organizational patterns according to the values of AT-skew or AT-skew / GC-skew in their leading strands (Fig. 1C). As expected, most high-GC sequences belong to the typical pattern and most low-GC sequences belong to the atypical pattern (Fig. 1C). GC-skew, AT-skew, GS-bias, and GC-content also show strong phylogenetic signals which can be reflected by the high values of Pagel’s λ (Fig. 2). Correspondingly, species possessing atypical sequences (i.e., with positive AT-skew in the leading strand) are highly clustered in the phylogenetic tree (Fig. 2). This implies that, high values of AT-skew in the leading strand of these species might be solely owing to shared evolutionary history ^39^. Hence, we applied phylogenetically independent contrasts (PIC) to explore the associations among them in two patterns separately. Results indicate that in the case of typical patterns, unlike the correlation pattern described above there is a strong negative correlation between AT-skew and GC-skew with a coefficient of -0.57 (adjusted p-value < 0.05, Fig. 3A). This pattern agrees with the previous asymmetrical deamination hypothesis ^10^. Whereas in the case of atypical pattern, we detected a positive correlation between AT-skew and GC-skew with a coefficient of -0.30 (adjusted p-value < 0.05, Fig. 3B). These results reveal a robust difference between two organizational patterns not only by the change in the values of AT-skew but also by the shift of correlation patterns of these organizational features, which suggests the participation of a different molecular mechanism likely promoted by PolC as previously assumed ^17^. That is, prokaryotic genomes inherently prefer an atypical organizational pattern with jointly increased GC-skew, AT-skew and GS-bias in the leading strand when have evolved to low GC-content (Fig. 3B).

**Figure 2.**
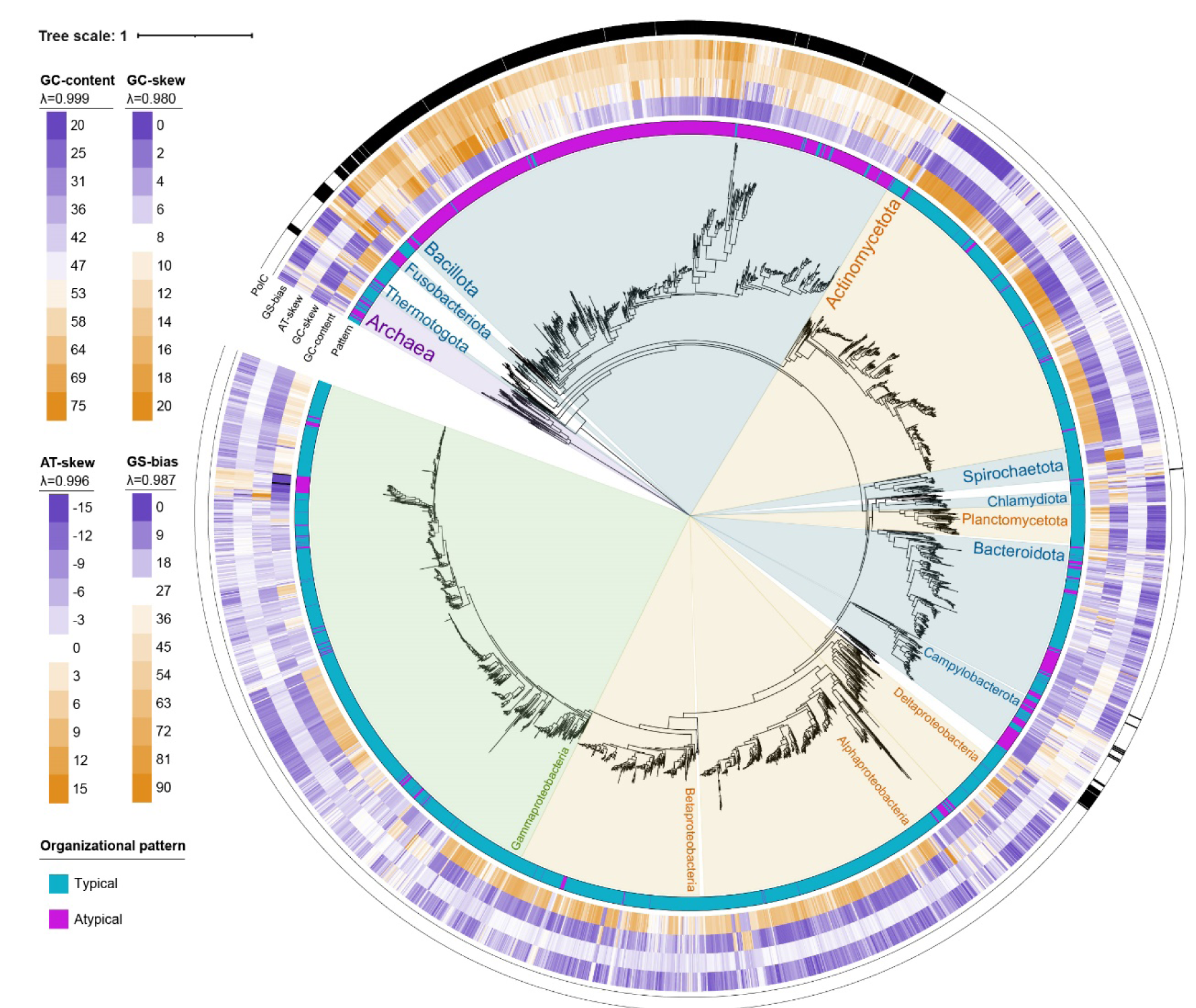
Distributions of GC-content, GC-skew, AT-skew, GS-bias and PolC on the tree of life of prokaryotes. Typical and atypical organizational patterns were labeled with different colors in the most inner circle. A maximum likelihood phylogenetic tree of 3329 representative genomes was constructed from the concatenated alignment of 27 ribosomal proteins and 3 RNA polymerases using the LG+R10 substitution model. Values of Pagel’s λ for each feature are labeled above the legend.

**Figure 3.**
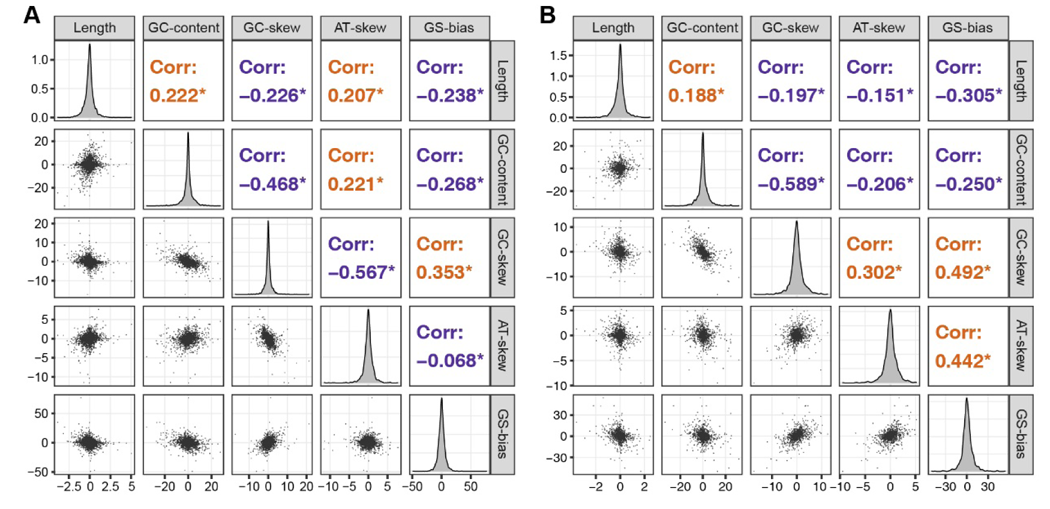
Two distinct patterns of prokaryotic genome organization identified based on the residuals from phylogenetically independent contrasts. (A) Pairwise correlation coefficients between the residuals of GC-content, GC-skew, AT-skew, GS-bias, and sequence length in typical pattern (i.e., the pattern where the leading strand displays AT-skew -). (B) Pairwise correlation coefficients between the residuals of GC-content, GC-skew, AT-skew, GS-bias, and sequence length in atypical pattern (i.e., the pattern where the leading strand displays AT-skew +). Asterisks indicate adjusted p-value < 0.05.

### Mutation and protein-coding genes determine the organizational pattern

Identifying these correlations and organizational patterns is insufficient for inferring causal mechanisms, especially distinguishing between the effects of mutational biases and selection which has long been controversial ^19,40,41^. Normally mutational biases have been thought to globally affect the genome sequences ^42^. Therefore, we further analyzed the uniformity of GC-skew and AT-skew across the leading strands. It can be found that the nucleotide skews display strong agreements between encoding (i.e., the coding genes on leading strands) and noncoding strands, with correlation coefficients 0.77 and 0.63 for GC-skew and AT-skew, respectively (Fig. 4A & B). Surprisingly, the GC-skew and AT-skew of anticoding strands (i.e., the complementary sequences of genes on the lagging strand) both show strong heterogeneity, negatively correlated with those of encoding and noncoding strands (Fig. 4A & B). The findings hint at the plausible influence of mutational biases, indicated by the global nucleotide skews of the leading or lagging strands, and distinctive mechanisms underlying the nucleotide skews of protein-coding genes.

**Figure 4.**
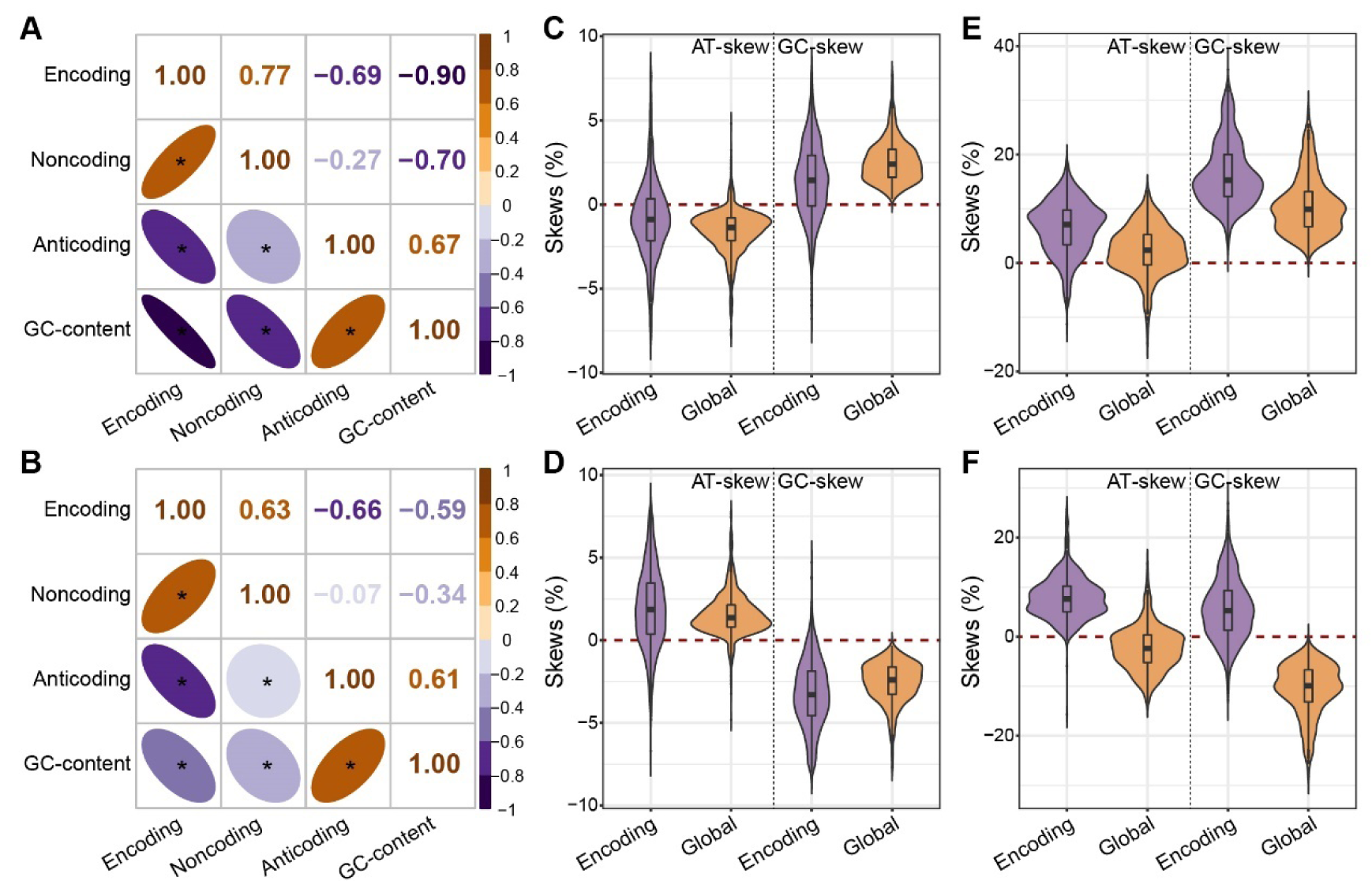
Relationships between the nucleotide disparities of coding genes and that of global strand. (A) Pairwise correlation coefficients between GC-content and GC-skew of encoding strand, anticoding strand, and noncoding strand. (B) Pairwise correlation coefficients between GC-content and AT-skew of encoding strand, anticoding strand, and noncoding strand. (C) Comparison of global nucleotide skews and the nucleotide skews of encoding sequences in the leading strand of high-GC (GC-content >= 55%) genomes. (D) Comparison of global nucleotide skews and the nucleotide skews of encoding sequences in the lagging strand of high-GC genomes. (E) Comparison of global nucleotide skews and the nucleotide skews of encoding sequences in the leading strand of low-GC (GC-content <= 45%) genomes. (F) Comparison of global nucleotide skews and the nucleotide skews of encoding sequences in the lagging strand of low-GC genomes.

Notably, GC-skew and AT-skew of both encoding and anticoding strands are highly correlated with GC-content (Fig. 4A & B). Indeed, regardless of whether it is on the leading or the lagging strand, the encoding strands have consistent nucleotide skews with the global strand in high-GC genomes (Fig. 4C & D). Separate nucleotide usages between the coding genes on the leading strand, mostly with positive GC-skew and negative AT-skew, and that on the lagging strand, mostly with negative GC-skew and positive AT-skew, are likely a result of the adaption to mutation context ^25,43^. Whereas, in low-GC genomes regardless of whether it is on the leading or the lagging strand, the encoding strands mostly have positive values of GC-skew and AT-skew, leading to a discordance in nucleotide skews on the lagging strand (Fig. 4E & F). Whatever the mutational pattern is, according to the Watson-Crick pairing rules only one strand, the leading strand because of the presence of essential and highly expressed genes ^5^, can reach an agreement between the GC-skew and AT-skew of protein-coding genes and global nucleotide skews (Fig. 4E & F). As a result, an atypical mutational pattern may have evolved to guarantee both GC-skew + and AT-skew + in the leading strand to protect essential genes ^17^. In parallel, lots of genes on the lagging strand should have been driven to the leading strand to avoid a mass of deleterious mutations caused by an uncoordinated mutational pattern, which can be verified by a large proportion of the explained variance in GS-bias by the nucleotide skews in lagging strands (Fig. S4). Collectively, results suggest that both mutational biases and the nucleotide skews of protein-coding genes contribute to a change from the typical to atypical organizational pattern in low-GC genomes under natural selection.

### Natural selection and genetic code determine the skew pattern of genes

As shown above, negative correlations between the nucleotide skews of coding genes and GC-content hold the key for the dramatic shift of organizational patterns (Fig. S5). Actually, four nucleotides are not equally used by the codons of different amino acids in standard genetic code (Fig. 5A), possibly due to multiple optimizations ^44^. Further, 20 amino acids are not equally used either (Fig. 5B), with the relative proportions constrained by natural selection though being highly variable ^45^. Our previous study has demonstrated that the usages of amino acids and synonymous codons both were powerfully affected by the genomic GC-content ^33^. Therefore, we conjecture here that the increased nucleotide skew driven by reduced GC-content derives from constrained amino-acid usage together with the special configuration of genetic code.

**Figure 5.**
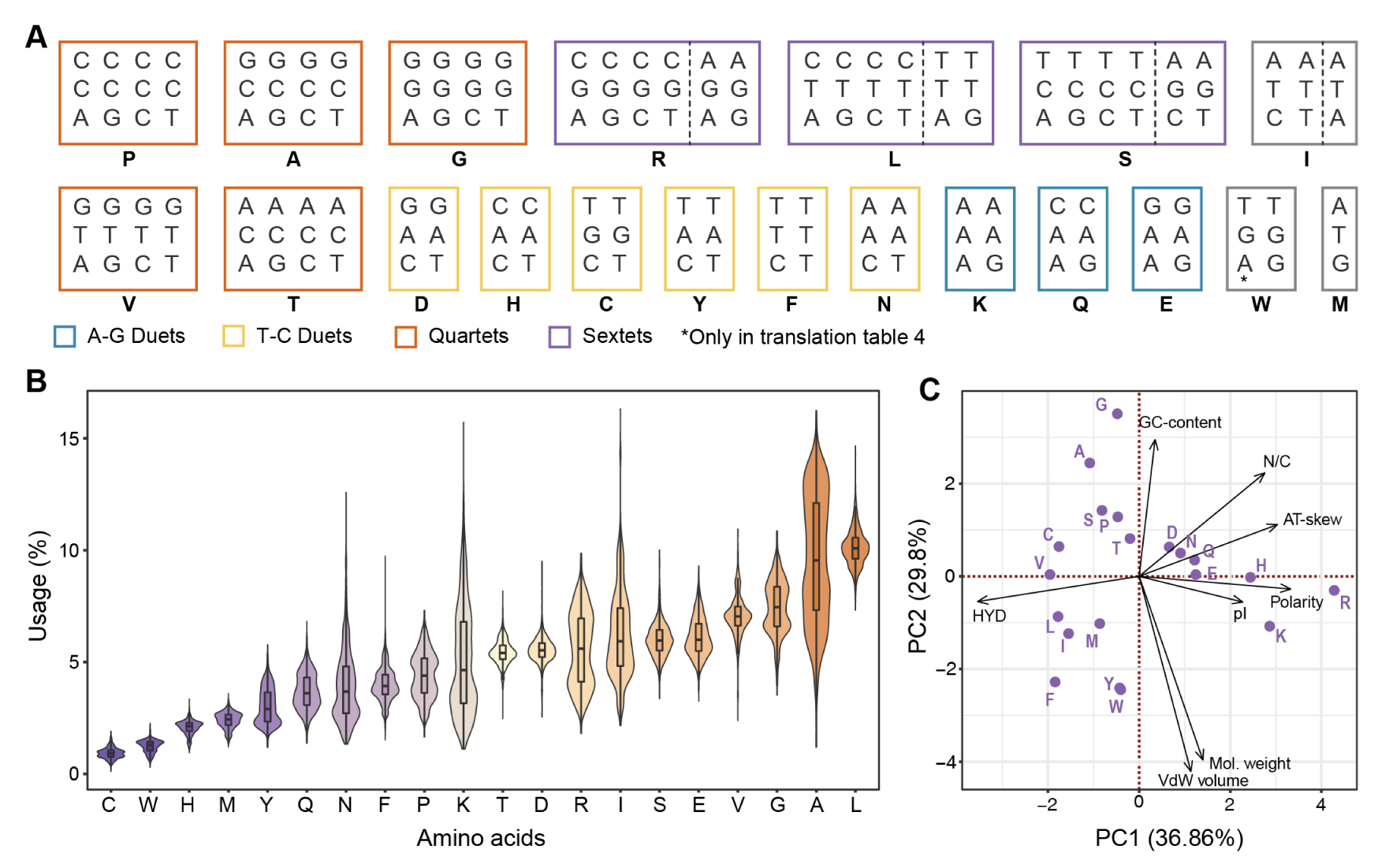
Genetic code and amino-acid usage of prokaryotes. (A) Configuration of the genetic code used by prokaryotes. A distinction is made between sextets, quartets, T-C duets, and A-G duets. (B) Variation in the proportion of 20 proteinogenic amino acids. (C) Principal component analysis of the physicochemical properties and nucleotide composition of 20 proteinogenic amino acids. The influences of different properties including molecular weight, VdW (Van der Waals) volume, hydrophobicity (HYD), isoelectric point (pI), polarity and N/C ratio, and nucleotide composition of codons including GC-content and AT-skew are denoted by arrows.

To verify our conjecture, we established three neutral models with amino-acid usages unrestricted for genomes with high GC-content (65%), middle GC-content (50%) and low GC-content (35%), respectively (see Methods). Meanwhile, three corresponding natural models with amino-acid usages satisfying the constraints of selection (that is, the proportions of all amino acids lay in the 95% confidence interval estimated using real sequences, see Methods) were also established (Fig. 6A, B & C). Results suggest that in neutral models GC-skew gradually goes up with reduced GC-content, whereas AT-skew displays no deviations (Fig. S6). This implies that the effect of GC-content (on the usages of amino acids and synonymous codons) alone is not sufficient to drive the AT-skew. However, in natural models, both GC-skew and AT-skew gradually rise with reduced GC-content (Fig. 6A, B & C). Positive GC-skew and AT-skew of coding genes in low-GC genomes appear to be an inevitable consequence of the joint impact of genetic code and natural selection, supporting our conjecture.

**Figure 6.**
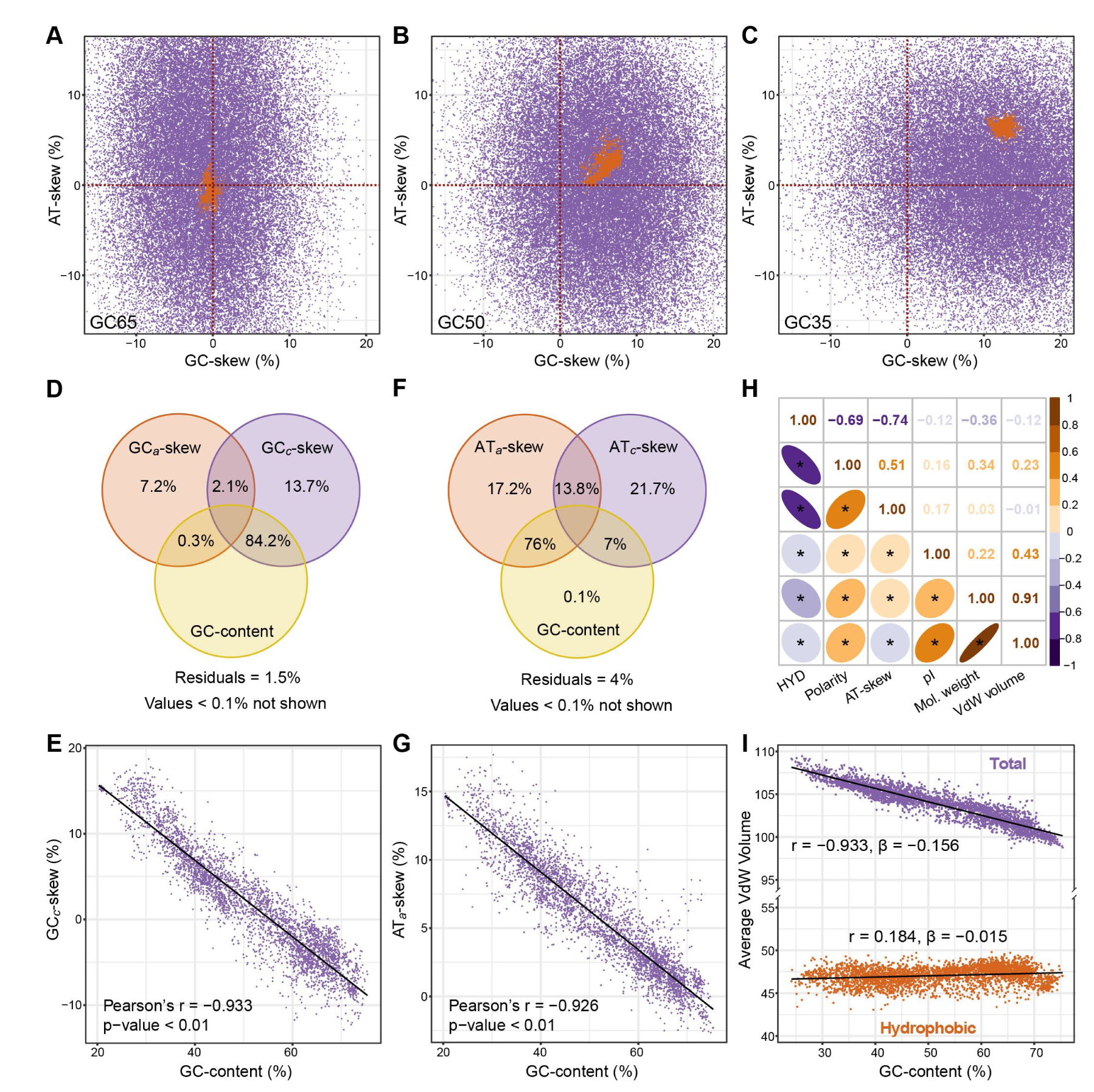
Mechanism for the evolution of nucleotide skews of protein-coding genes. (A) Variations of GC-skew and AT-skew in neutral (shown in violet color) and natural (shown in orange color) models with high GC-content (i.e., 65%). (B) Variations of GC-skew and AT-skew in neutral and natural models with middle GC-content (i.e., 50%). (C) Variations of GC-skew and AT-skew in neutral and natural models with low GC-content (i.e., 35%). (D) Variance partitioning analysis showing the fractions of variance in GC-skew explained by GC*_a_*-skew (i.e., GC-skew contributed by the usage of amino acids only), GC*_c_*-skew (i.e., GC-skew contributed by the usage of codons only), and GC-content. (E) Correlation plot showing the strong agreement between GC*_c_*-skew and GC-content (F) Variance partitioning analysis showing the fractions of variance in AT-skew explained by AT*_a_*-skew (similar to GC*_a_*-skew), AT*_c_*-skew, and GC-content. (G) Correlation plot showing the strong agreement between AT*_a_*-skew and GC-content. (H) Pairwise correlation coefficients between AT-skew and average properties of amino acids in neutral GC35 model. (I) Comparison of the relationship between average VdW volume of total amino acids and GC-content, and that between the fraction contributed by hydrophobic amino acids (i.e., the hydrophobic bulk) and GC-content, where values of *β* represent the slope of linear regression equations. Solid lines in E, G and I represent fits of linear regression.

To clarify the mechanism of such impact, we performed variance partitioning analysis of GC-skew and AT-skew. Results reveal that ∼84% of the variance in GC-skew is explained by the covariance between GC*_c_*-skew (i.e., GC-skew contributed by codon usage only) and genomic GC-content (Fig. 6D). GC-content and GC*_c_*-skew are indeed negatively correlated (with the coefficient of -0.933, or -0.933 in phylogenetically independent analysis), indicating that genomes have low values of GC-content also have biased codon usages resulting in high GC-skew (Fig. 6E & Fig. S7). This is probably attributed to a higher number of C_3_ synonymous codons (codons with C in their third position) compared with that of G_3_ synonymous codons (16 vs 13, Fig. 5A). Particularly, both the number (6 vs 3, Fig. 5A) and proportion of T-C duets are always higher than those of A-G duets (Fig. S8). When genome evolves towards lower GC-content, the transitions of C-to-T and G-to-A in the third position of these duets will lead to more loss of C than G thus causing a rise in GC-skew (Fig. S9).

Theoretically, such variation will also lead to a decline of AT-skew. Nevertheless, the existence of sextets, each of which can be considered as the combination of a quartet and a duet, complicates the situation (Fig. 5A). Evolving towards lower GC-content brings changes of arginine (R) and leucine (L) from using high-GC codons (R: CGC and CGG, L: CTC and CTG) to using low-GC codons (R: AGA, L: TTA), resulting in a net loss of C and a net gain of A (Fig. S10). Hence, the AT*_c_*-skew decreases initially and then increases with the decrease of GC-content (Fig. S11). In contrast, AT*_a_*-skew (i.e., AT-skew contributed by amino acid usage only) and GC-content together explain the majority (∼76%) of variance in AT-skew (Fig. 6F). Correspondingly, GC-content and AT*_a_*-skew are strongly negatively correlated, with the coefficient of -0.926, or -0.782 in phylogenetically independent analysis (Fig. 6G & Fig. S7), implicating that A-rich (amino acids encoded by codons with A content ≥ 50%, i.e., K, N, E and Q) instead of T-rich (amino acids encoded by codons with T content ≥ 50%, i.e., F, Y, C and L) amino acids are preferentially used in the case of low GC-content. Of these amino acids, both the A-rich amino acids K and N, and the T-rich amino acids F and Y increase their proportion with the decrease of GC-content due to low GC-content of their codons, however, the former more rapidly than the latter (Fig. S12). Taking the most A-rich amino acid K and the most T-rich amino acid F as an example, the usage of K grows more rapidly than that of F with the decrease of GC-content and ends up with higher proportion though starting with lower proportion (Fig. S13). Variation in the usage of these two amino acids alone explains more than 91% of the variance in AT*_a_*-skew, suggesting that the amino-acid usage likely shaped by natural selection deeply influences AT-skew (Fig. S14).

### Hydrophobic bulk conservation determines the amino acid usage of genes

The above results raise a key question: how does selection influence the amino-acid usage, and thereby AT-skew? When considering the amino acid properties presumably constrained by natural selection, several interesting findings emerge. Firstly, amino acids encoded by codons with lower GC-content have higher molecular weight, VdW (Van der Waals) volume and N/C ratio (Fig. 5C & Supplementary data 3), consistent with the results that these three properties all display strong relevance to GC-content among different neutral models (Fig. S15). This implies that evolving towards lower GC-content will also bring substitutions from amino acids with small volumes to ones with large volumes. Secondly, amino acids encoded by codons with higher AT-skew have lower hydrophobicity and higher polarity (Fig. 5C & Supplementary data 3). The hydrophobicity (hydropathy index) takes into account not only the property of amino acid residues itself but also their position in protein (being on the surface or buried inside) ^46^. According to current knowledge, when a protein folds hydrophobic amino acids generally get buried inside to form a hydrophobic core to stabilize the structure, thereby protein is quite sensitive to the mutation of buried hydrophobic amino acids ^47^. Amino acid substitution thus occurs more easily on the surface in contrast to protein interior, especially that with a large volume change ^48,49^. It is reasonable to infer that when genomes evolving towards lower GC-content, the substitution of amino acids on the protein surface, that is, to ones with low hydrophobicity (and with high AT-skew of codons) should occur more frequently than in the protein interior (to ones with high hydrophobicity), which agrees perfectly with the finding of the comparison between the K and F (Fig. S13 & Supplementary data 3).

Despite the positive correlation between GC-content and average hydrophobicity of amino acids, low GC-content by itself is insufficient to induce low hydrophobicity as shown in our established neutral models, suggesting that variation in hydrophobicity results from natural selection (Fig. S15 & Fig. S16). Correspondingly, in the GC35 (GC-content 35%) neutral model AT-skew is particularly correlated to hydrophobicity with a correlation coefficient of -0.74, indicating that high AT-skew also results from such selection supporting our conjecture (Fig. 6H). Of the most hydrophobic amino acids (i.e., I, L, V, F, C, M, A and G) which are largely buried inside protein ^46^, A and G with small VdW volume decrease their proportion with the decline of GC-content while I and F with large VdW volume increase their proportion (Fig. S17). As a result, albeit with the decrease of the total usage of these hydrophobic amino acids (Fig. S17), the hydrophobic fraction of average VdW volume (average VdW volume contributed by eight hydrophobic amino acids, i.e., hydrophobic bulk, see methods) is weakly influenced by GC-content (r = 0.184, β = -0.015), in contrast with the average VdW volume of total amino acids which is strongly influenced (r = -0.933, β = -0.156, Fig 6H). Such conservation of hydrophobic bulk likely results from maintaining stable structure of proteins under selection, and fundamentally determines the amino acid usage and AT-skew of genes.

## Discussion

In prokaryotic genomes, GC-content, GC-skew, AT-skew, and GS-bias are essential compositional and organizational features, and a growing body of evidence suggests tight associations between them ^5,14–16,24,25,35,50,51^. Although both mutational biases and selection have been proposed to account for the origins, their dramatic variations are not yet satisfactorily explained. Here, we have performed more systematical analysis of bulk sequence data contributing to a novel mechanistic framework, centered around the variation in GC-content which has been shown to result from mutational biases in ancient adaptive processes ^33^, for understanding the evolution of genome organization (Fig. 7). Firstly, both GC-skew and AT-skew increase with decreasing GC-content in protein-coding genes, mostly determined by synonymous substitutions because of the special configuration of genetic code, and by nonsynonymous substitutions owing to hydrophobic bulk conservation, respectively. Secondly, such variations in nucleotide skews of protein-coding genes interact with mutational biases leading to two distinct organizational patterns: (i) Typical pattern mainly in high-GC genomes with putative G- and T-biased mutational pattern in the leading strand and minor values in GC-skew, AT-skew, and GS-bias, and (ii) atypical pattern mainly in low-GC genomes with G- and A-biased mutational pattern in the leading strand and high values in GC-skew, AT-skew, and GS-bias (Fig. 1 & Fig. 7). As described in previous studies, the shift in organizational pattern is likely to have been achieved through horizontal gene transfer and the evolution of PolC ^14,16,17,19^. Although ancient evolutionary processes can be neither easily validated nor easily invalidated via experiments ^33^, parallel emergence of PolC and atypical AT-skew in multiple clades including Bacillota, Fusobacteriota and Campylobacterota further supports the above hypothesis (Fig. 2).

**Figure 7.**
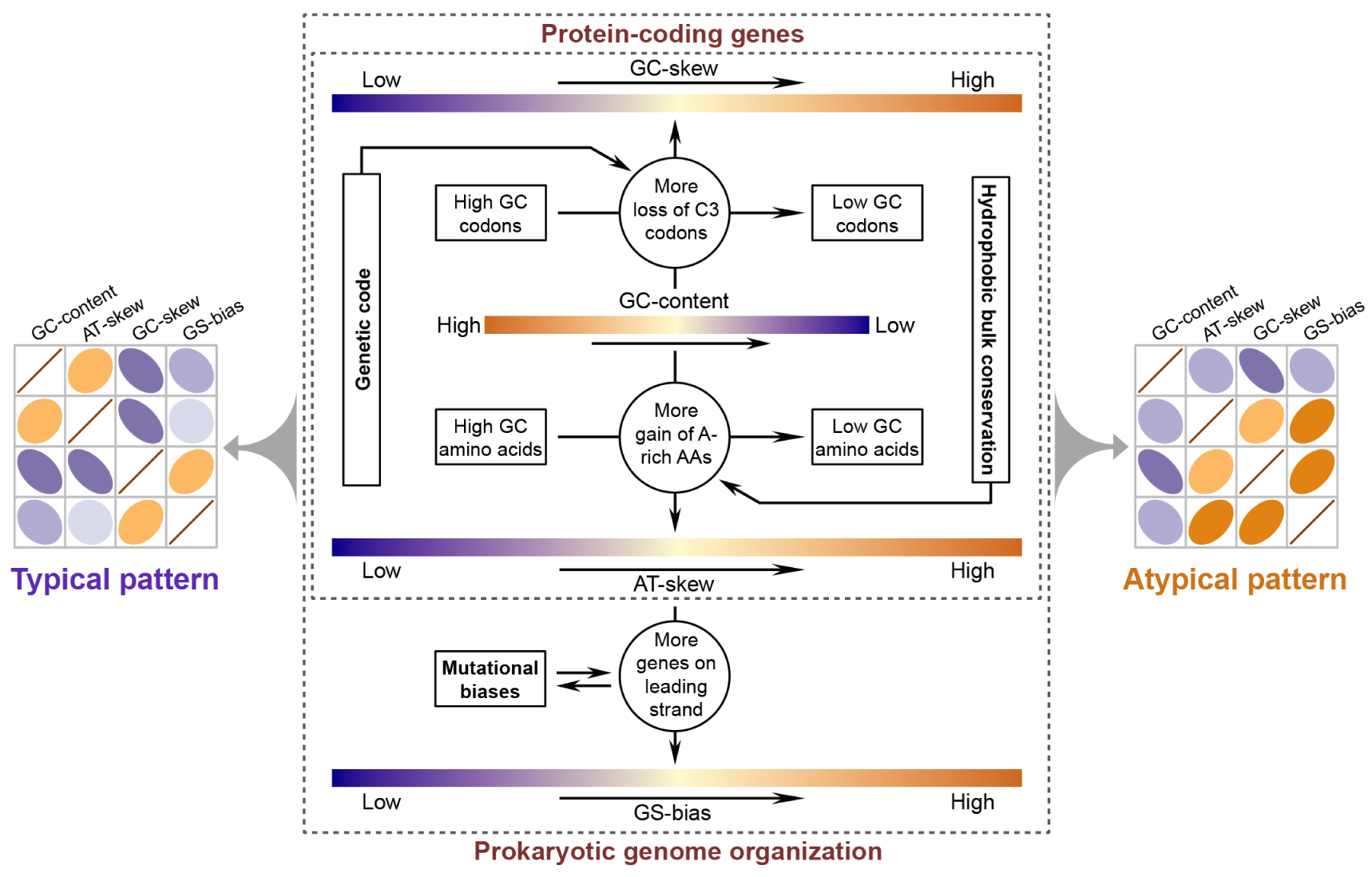
Evolutionary mechanism for the organization pattern of prokaryotic genome. Arrows indicate the change trend of GC-content, GC-skew, AT-skew, and GS-bias as well as the causal structure generating associations between them. Pairwise diagrams at the left and the right side indicate different correlation patterns which are same as those displayed in Figure 3.

In our mechanistic framework, negative correlations between GC-content and GC- & AT-skew of protein-coding genes are a critical nexus for genome organization (Fig. 7). Biases in amino-acid usage and codon usage leading to these nucleotide skews have been attributed to transcription-associated selection for the optimization to nucleotide and tRNA pools ^28,52^. However, this effect resides mainly in highly expressed genes accounting for only a small fraction of the overall coding sequences ^53^. Other views concerning energy and resource efficiency argue that selection for cheaper nucleotides and amino acids drive nucleotide skews, especially the atypical AT-skew, and gene distribution ^14,19,27^. Nevertheless, at least two facts contradict such hypotheses: (i) The availability of A is much higher than that of “cheaper” T, due to the metabolic role of ATP as an energy currency ^54^, and (ii) most microbes, like heterotrophic bacteria, have no need for de novo synthesis of all 20 amino acids during growth, which implies that the cost of an amino acid can be different among different species and under different environmental conditions ^55,56^. Selection on energy and resource efficiency should be vulnerable to ever-changing environments over long timescales, particular in bacteria known for rapid adaptive evolution ^57^, in conflict with strong phylogenetic signals of organizational features (Fig. 2). Moreover, it has also been demonstrated recently that genetic code is not, as declared by the resource efficiency hypothesis, optimized for resource conservation ^44,58^. In contrast, both hydrophobicity and molecular volume are crucial for amino acids with respect to their roles in the protein, which also manifests in the optimization of genetic code ^59–61^. While substitutions are more likely to occur between amino acids with similar chemical properties, those from small hydrophobic ones to large hydrophobic ones should be quite scarce to keep proteins hydrosoluble and in a compact form ^49,62^. According to the findings in our study, we conjecture that high hydrophobic bulks of T-rich amino acids, which may influence protein structure stabilization, instead of high energy costs account for the negative correlation between GC-content and AT-skew (Fig. 7).

Collectively our results suggest that mutational biases provide the driving force, and genetic code and hydrophobic bulk conservation determine the organization pattern, which is reflected by the value of AT-skew as well as its correlations with GC-content and other features. The effects of mutational biases on genome evolution likely have been underestimated in previous studies, mainly owing to the suspicion of neutral processes in prokaryotes ^31,63^. However, mutational biases can also contribute much during adaptive processes, and even serve as an essential mechanism of adaptation by generating adaptive mutations ^34,64–67^. The relationship between the mutational biases driving GC-content variation and that giving rise to nucleotide skews, as well as the molecular role that PolC has played, of course, require further exploration. Still, this study again highlights the importance of evolutionary processes operating at “lower levels of organization” as Lynch stated ^32^, including the configuration of genetic code, protein folding, and mutational biases likely imposed by DNA replication and repair. Accidental by-products generated indirectly in these evolutionary processes might confuse the role of natural selection, cautioning against directly linking every aspect of genome to an environmental interpretation.

## Methods

### Sequence data collection and filtration

A total of 4,218 NCBI-defined representative genomes (assembly level: complete and chromosome) were downloaded on 30 March 2021 from the NCBI RefSeq database (https://www.ncbi.nlm.nih.gov/refseq/). Of these, 3,984 and 234 genomes belong to bacteria and archaea, respectively (Supplementary data 1). NCBI-annotated complete sequences (including chromosomes and plasmids) with sequence length of more than 200 kb were selected for further processing. Cumulative curves of GC-skew, AT-skew, and GS-bias (cumulative gene distribution, where 1 represents a gene on the leading strand and -1 represents a gene on the lagging strand) were plotted based on a sliding window approach using custom Python and R scripts. The skew index of GC-skew (GCskewI) and AT-skew (ATskewI) were also calculated using scripts from previous study ^36^. Sequences with GCskewI or ATskewI less than 0.5 were manually inspected according to their Z curves to filtrate ones without apparent skew pattern. Eventually, a total of 4,012 complete sequences were obtained (Supplementary data 2), and the leading strand for each sequence (i.e., sequence region from the origin to the terminus of replication, which corresponds to that from the minimum to the maximum in the GC-skew curve) was truncated using Seqtk (https://github.com/lh3/seqtk/). Sequence data and Z curves of all obtained leading strands have been deposited in the Figshare repository (http://dx.doi.org/10.6084/m9.figshare.24314023).

### Quantitative analysis of organizational features

The sequence length, GC-content, GC-skew, AT-skew, and GS-bias (Gene strand bias) of all leading strands were calculated using custom Python and Perl scripts. Prodigal v2.6.3 was used to identify protein-coding genes in all sequences ^68^. The GS-bias was calculated as:

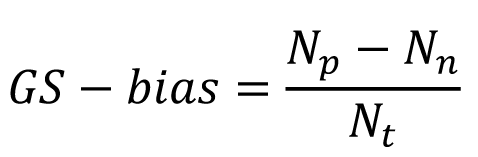

Where *N_p_*, *N_n_* and *N_t_* indicate the number of genes on the leading strand, the number of genes on the lagging strand and the total number of genes, respectively. According to the position of genes, the sequences of encoding regions (i.e., where coding genes lie), anticoding regions (i.e., where the complementary strand of coding genes lie) and the sequences of noncoding regions were identified and extracted from the leading strands. GC-content and organizational features of them were separately calculated to evaluate the intragenomic heterogeneity. GC-skew and AT-skew of coding genes from each leading strand were also calculated. Moreover, the nucleotide skew contributed by the amino acid usage only was calculated by fixing the synonymous codon usage

(taking the GC-skew as an example):

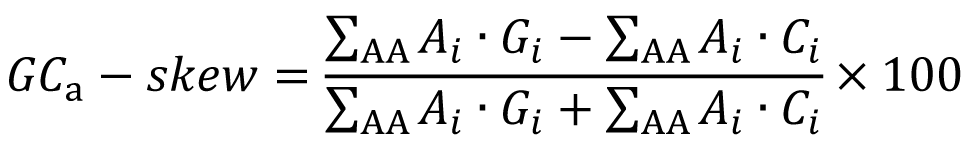

where *A_i_* is the abundance of an amino acid (AA), *G_i_* is the proportion of base G and *C_i_* is the proportion of base C in the codons of an amino acid. The subscript *i* (from 1 to 20) indicates the *i* th amino acid. The GC-skew contributed by codon usage only was calculated by fixing the amino acid usage:

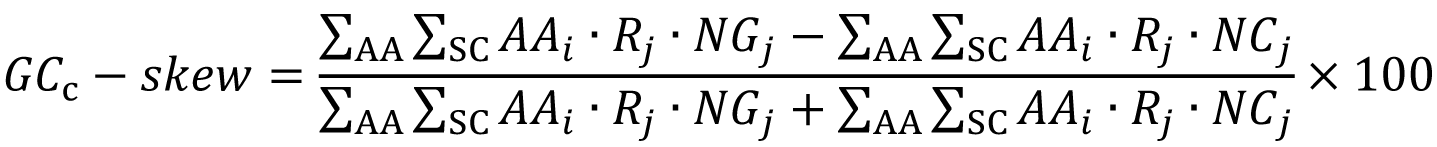

where *AA_i_* is the average abundance of an amino acid, *NG_j_* is number of base G and *NC_j_* is number of base C in a synonymous codon (SC) of a specific amino acid. *R_j_* is the relative abundance of a synonymous codon of a specific amino acid. The subscript *j* (from 1 to 2∼6) indicates the *j* th synonymous codon.

### Phylogenetic analysis of organizational features

For phylogenetic analysis, a concatenated alignment of ribosomal proteins and RNA polymerases was generated using the MarkerFinder pipeline ^69^. The alignment was further trimmed to remove poorly aligned positions using trimAl v1.4. rev22 with the -automated1 option ^70^. A maximum likelihood phylogenetic tree was then constructed from the trimmed alignment using IQ-TREE v2.2.0 with the LG+R10 substitution model and options -B 1000, --bnni, and -T AUTO ^39,71^. This tree was rooted using the divergence between archaea and bacteria and was visualized using iTOL ^72^. Pagel’s λ, a measure ranging from 0 to 1 and a high value of it represents a strong phylogenetic signal, was estimated for GC-content, GC-skew, AT-skew and GS-bias in the above phylogenetic tree using the phytools package in R ^73^. Furthermore, phylogenetically independent contrasts were performed for length, GC-content, GC-skew, AT-skew and GS-bias to test the correlation pattern of genome organization without phylogenetic influence using the ape package in R ^74^. A Hidden Markov Model (HMM) profile was downloaded from KOfam (ftp://ftp.genome.jp/pub/db/kofam/) to identify orthologues of PolC (K03763) using hmmsearch command (E-value ≤ 10^-5^; alignment coverage ≥ 0.7) in HMMER v3.1b2 ^75^. The phylogenetic distribution of PolC orthologues was then explored in the above phylogenetic tree.

### Construction of neutral and natural models

Evolutionary models of coding genes were constructed separately for genomes with high GC-content (65%), middle GC-content (50%), and low GC-content (35%). Of these, 3 neutral models were established with amino-acid usages unrestricted to test how GC-content influences GC-skew and AT-skew due to the configuration of genetic code. Taking that for the low GC-content (35%) as an example, the amino-acid usage and codon usage of protein-coding genes from genomes with low GC-contents (35 ± 2%) was first investigated. Both the average abundance of each amino acid and the average relative proportions of their synonymous codons were calculated. Next, the neutral model was established by generating 50,000 groups of random numbers using a custom Python script to represent 50,000 genomes evolved neutrally. Each group consists of 20 random numbers representing the proportions of 20 amino acids which satisfy the following requirements: (i) All the numbers range between the minimum and maximum of real amino-acid proportions in the above low GC-content genomes. 1. (ii) The sum of these numbers is equal to 1 (100%). (iii) The GC-content which can be computed according to the above calculated average relative proportions of codons is equal to 0.35 (35%), and (iv) the variance of these numbers is equal to the mean value of the variance of real amino-acid proportions. The natural model was established by generating 10,000 groups of random numbers which satisfy an additional requirement: Each number ranges between the 0.025 quantile and 0.975 quantile of real proportions of one corresponding amino acid. All evolutionary models and scripts are available on Figshare (http://dx.doi.org/10.6084/m9.figshare.24314023).

### Other statistical analyses and visualization

Other statistical analyses, including correlation analysis, regression analysis, and the calculation of means and variances were performed using R v4.1.1. Results generated in this study were mainly visualized using R package ggplot2. All estimated P values in multiple comparisons were adjusted in R using the Bonferroni method ^76^. Variance partitioning analysis (VPA) of the GC-skew and AT-skew were performed using R package vegan. The average properties of amino acids were calculated as described in our previous study ^33^. The hydrophobic bulk was calculated as:

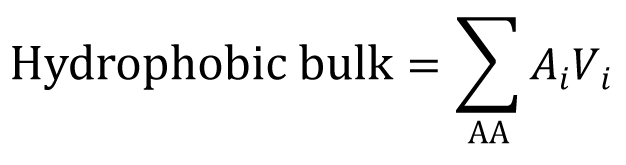

where *V_i_* is the VdW volume of an amino acid. The subscript *i* (from 1 to 8) indicates the *i* th amino acid from I, L, V, F, C, M, A and G. Circular diagrams of chromosome sequences from *Pseudomonas furukawaii* and *Shouchella miscanthi* were generated using Circos v 0.69 ^77^. Functional categories were assigned to each coding gene using BLAST+ against the latest COG database ^78^. Non-coding RNA genes were identified by searching the Rfam v14.5 database using Infernal v1.1.4 ^79^.

## Acknowledgments

We thank Guisong Chen from Guangdong *Magigene* Biotechnology Co., Ltd, for his assistance with the writing of Python scripts for the model construction, and Carolina A. Martínez-Gutierrez from Virginia Tech, for her suggestions helping to improve the manuscript. This work was supported by grants from the National Natural Science Foundation of China (Nos. 32370009 and 92351301).

## Notes

http://dx.doi.org/10.6084/m9.figshare.24314023

### Competing Interest Statement

The authors have declared no competing interest.

