## Supplemental Figure 1-17 for "Hydrophobic bulk conservation and genetic code determine the pattern of prokaryotic genome organization"

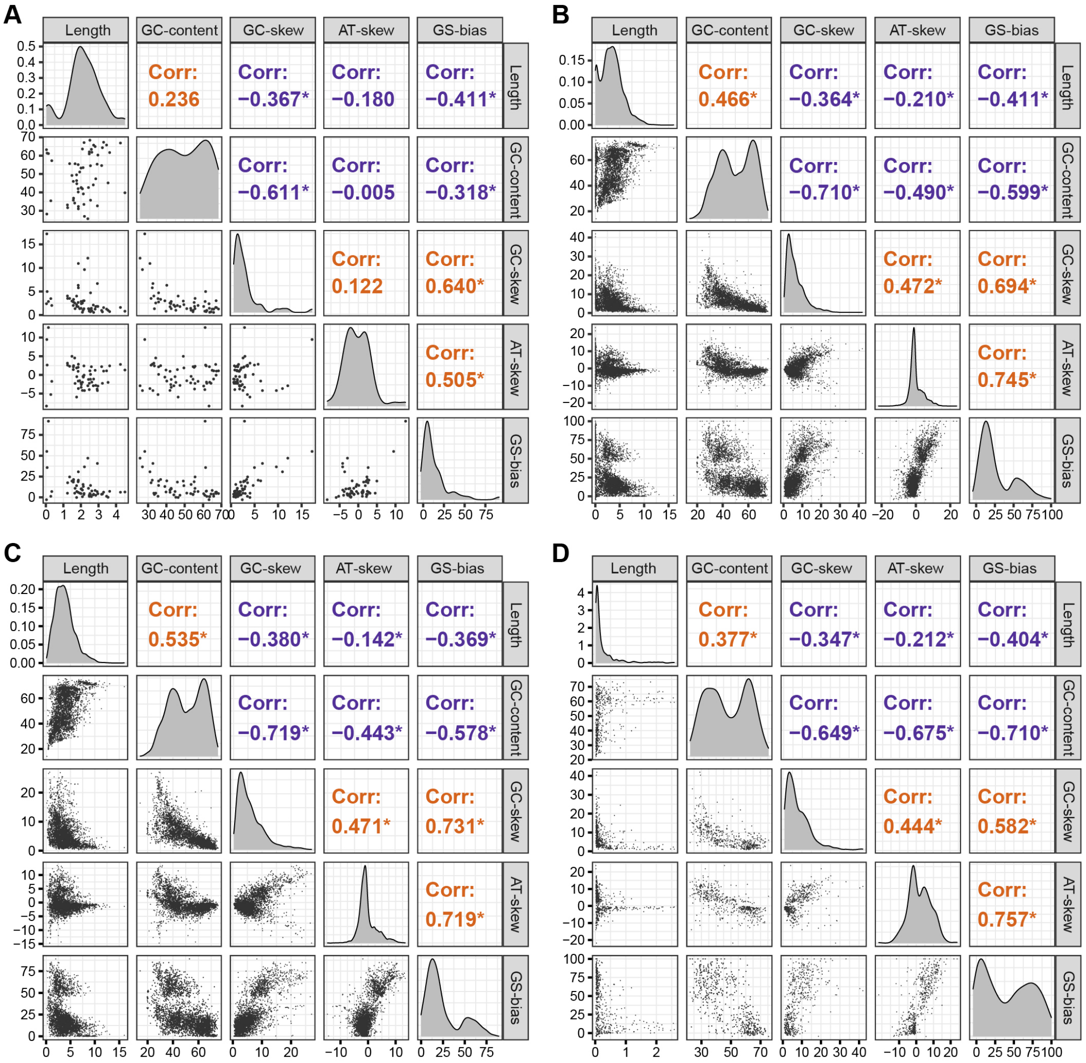


**Figure S1** Pairwise correlation coefficients between GC-content, GC-skew, AT-skew, GS-bias, and sequence length. (A) Correlation coefficients calculated using sequences from archaea. (B) Correlation coefficients calculated using sequences from bacteria. (C) Correlation coefficients calculated using chromosome sequences. (D) Correlation coefficients calculated using plasmid sequences.

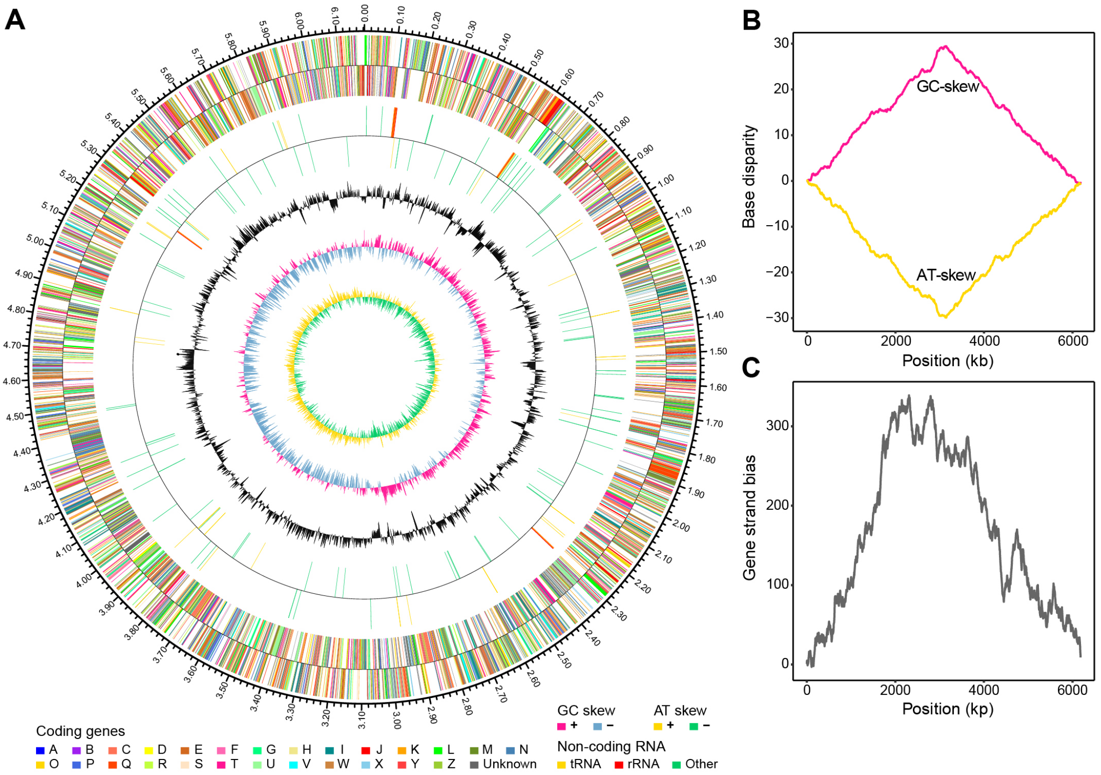


**Figure S2** Genome map of *Pseudomonas furukawaii*. (A) Circular representation of the chromosome sequence. From outside to the center: (i) coding genes on the forward strand (colored by COG categories); (ii) coding genes on the reverse strand (colored by COG categories); (iii) RNA genes on the forward strand; (iv) RNA genes on the reverse strand; (v) GC content; (vi) GC skew; (vii) AT skew. (A) Z curves of GC-skew and AT-skew. (C) Z curves of GS-bias.

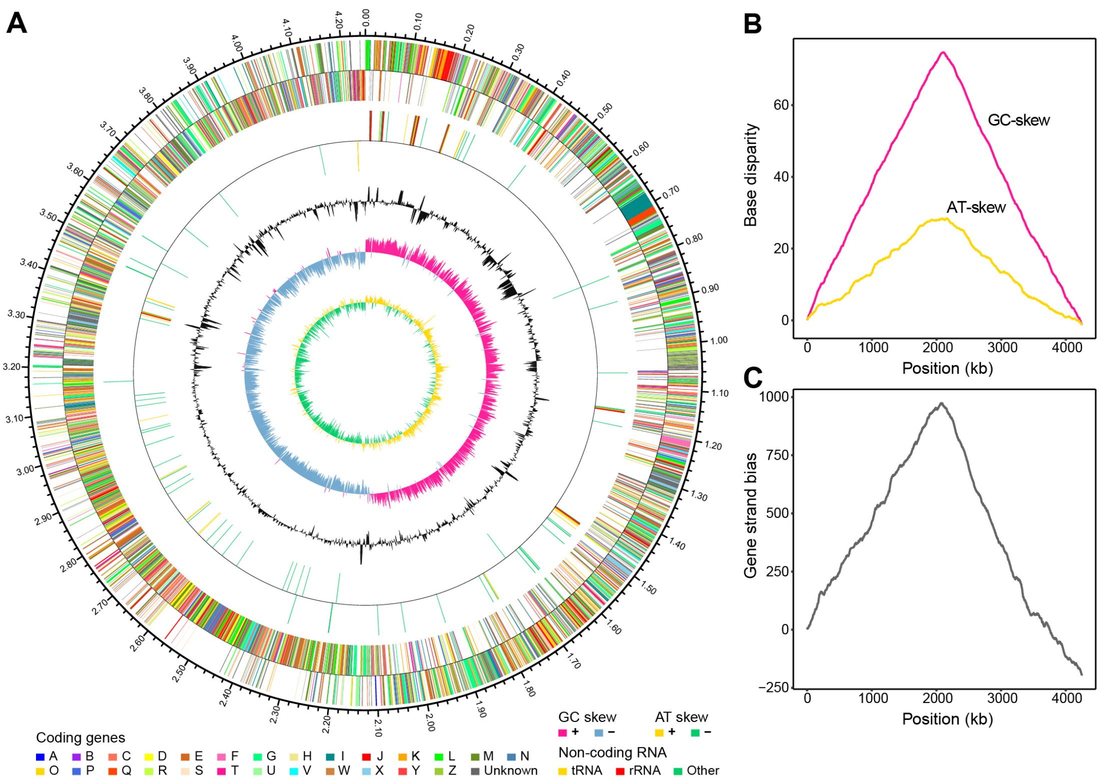


**Figure S3** Genome map of *Shouchella miscanthi*. (A) Circular representation of the chromosome sequence. From outside to the center: (i) coding genes on the forward strand (colored by COG categories); (ii) coding genes on the reverse strand (colored by COG categories); (iii) RNA genes on the forward strand; (iv) RNA genes on the reverse strand; (v) GC content; (vi) GC skew; (vii) AT skew. (A) Z curves of GC-skew and AT-skew. (C) Z curves of GS-bias.

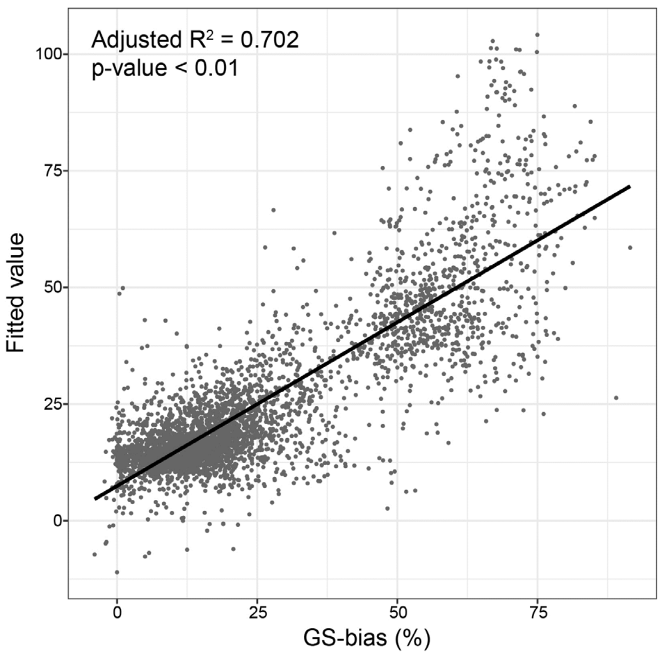


**Figure S4** Results of multiple regression analysis, where the GS-bias was explained by the global GC-skew of lagging strand, the global AT-skew of lagging strand, the GC-skew of coding genes on the lagging strand, and the AT-skew of coding genes on the lagging strand. Fitted values of GS-bias are shown in contrast to real values.

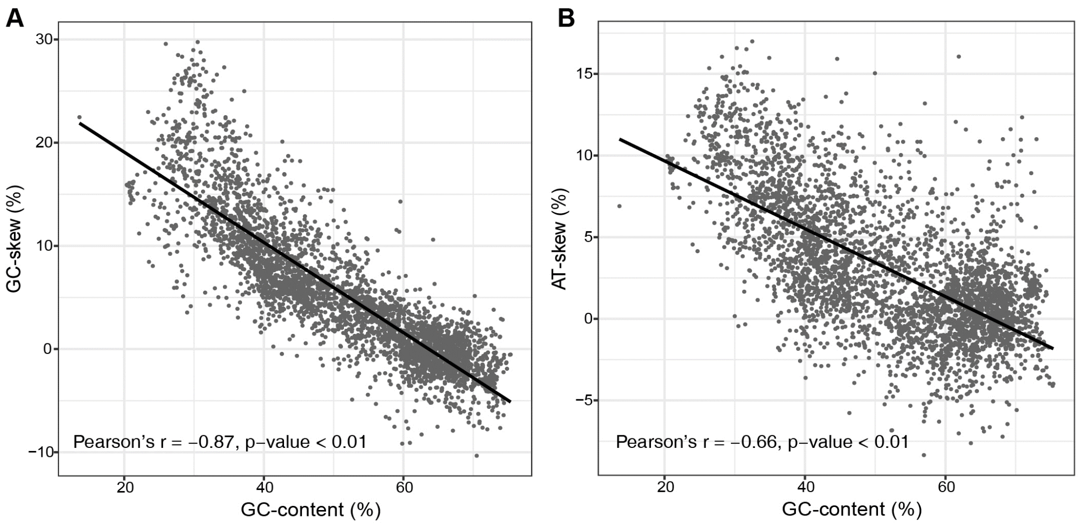


**Figure S5** Relationships between genomic GC-content and the nucleotide skews of coding genes. (A) Correlation plot showing high agreement between GC-content and the GC-skew of coding genes. (B) Correlation plot showing high agreement between GC-content and the AT-skew of coding genes. Solid lines in A and B represent fits of linear regression.

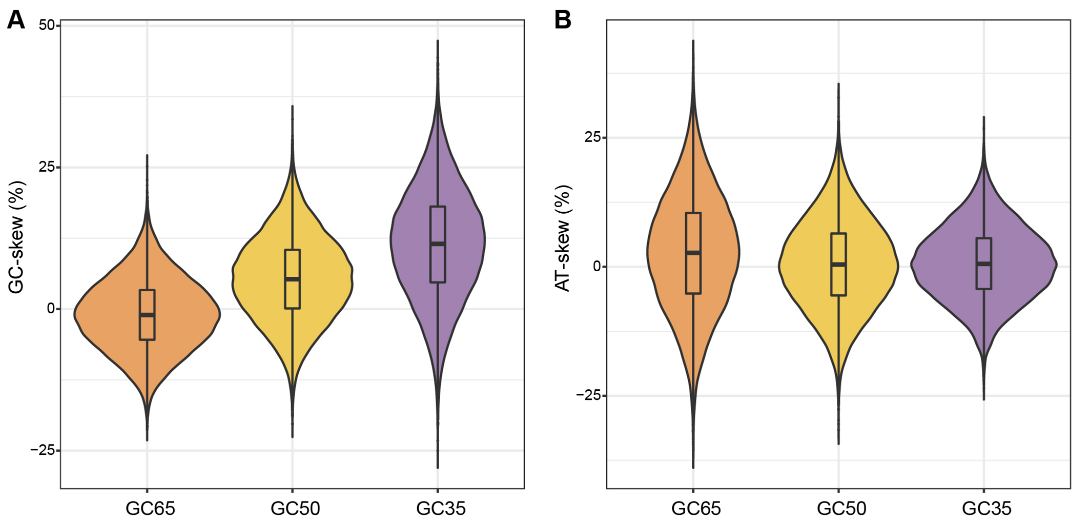


**Figure S6** Variations of GC-skew (A) and AT-skew (B) in established neutral modes with different levels of GC-content.

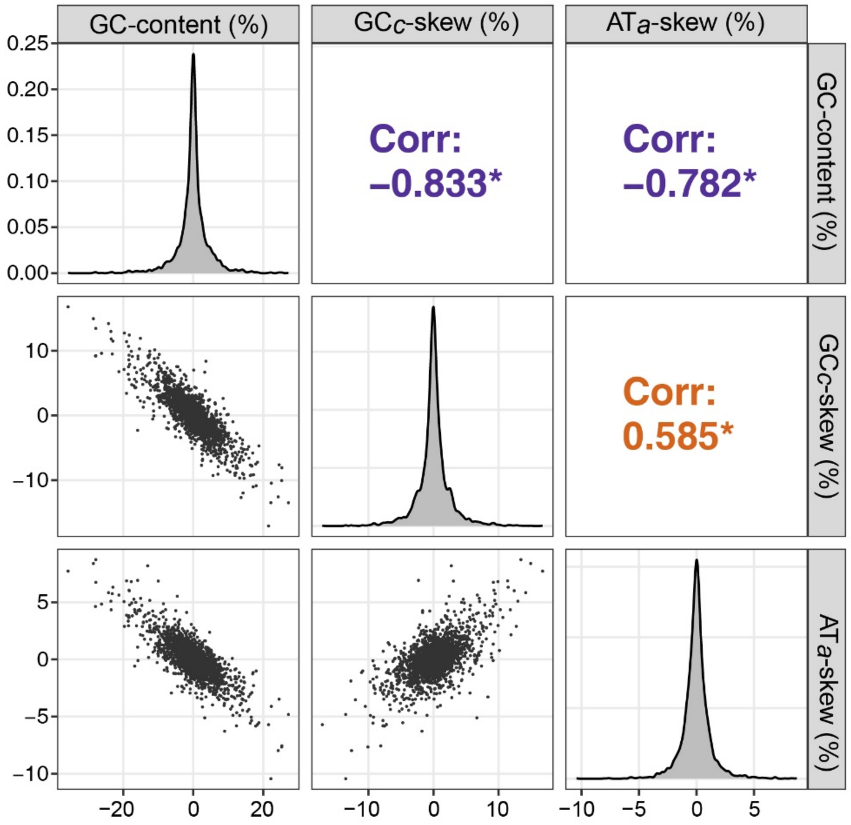


**Figure S7** Pairwise correlation coefficients between the residuals of $GC_c$-skew (i.e., GC-skew contributed by the usage of codons only), $AT_a$-skew (AT-skew contributed by the usage of amino acids only) and GC-content from phylogenetically independent analyses. Asterisks indicate adjusted p-value $< 0.05$.

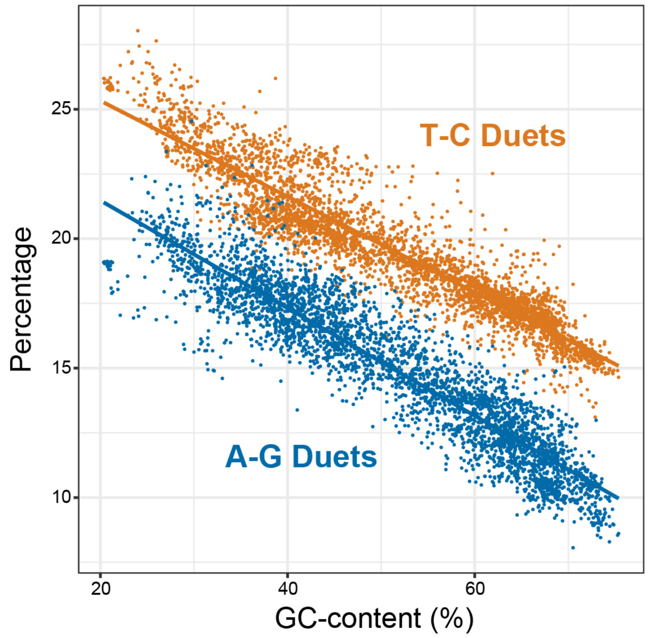


**Figure S8** Comparison between the usage of A-G duets (shown in blue) and T-C duets (shown in orange) with different values of GC-content.

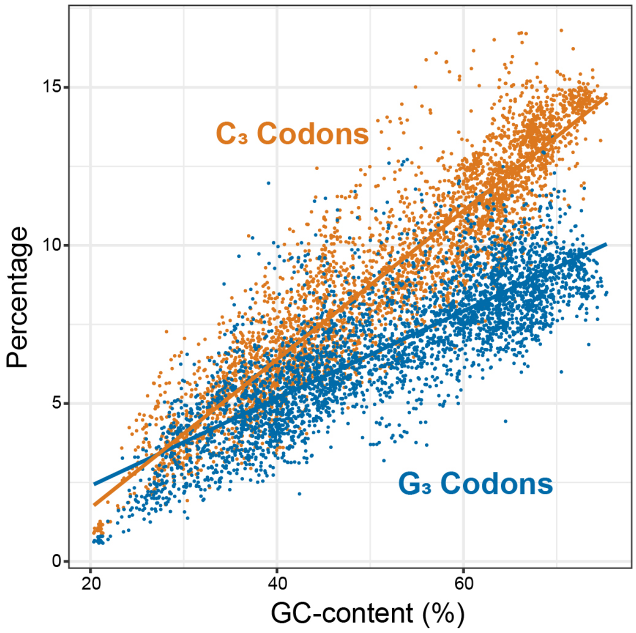


**Figure S9** Comparison between the usage of $G_3$ codons (codons with G in their third position) and C3 duets (codons with C in their third position) with different values of GC-content.

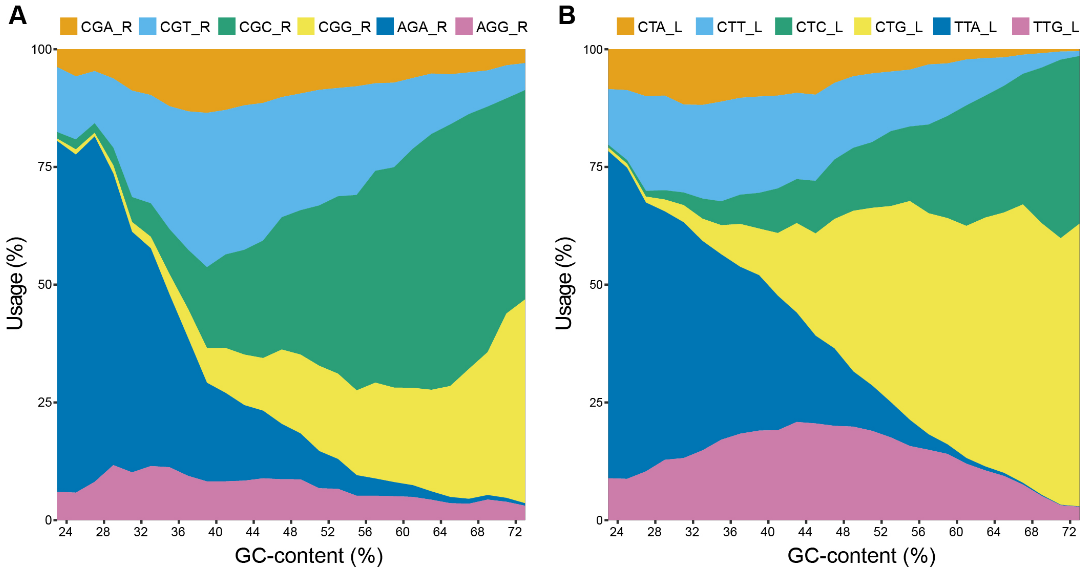


**Figure S10** Relative synonymous codon usage for sextets. (A) Relative synonymous codon usage for arginine (R). (B) Relative synonymous codon usage for leucine (L).

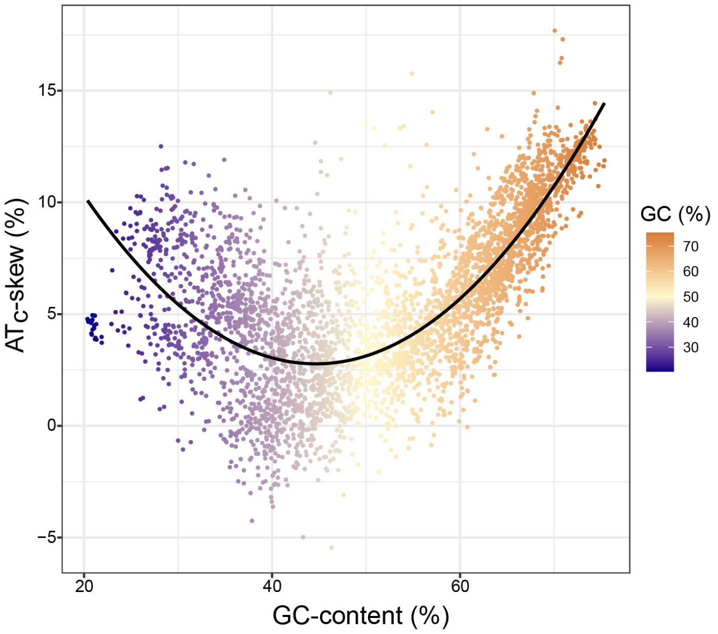


**Figure S11** Relationship between genomic GC-content and AT$_c$-skew (i.e., AT-skew contributed by synonymous codon usage only). Solid line in the plot represents the fit of polynomial regression.

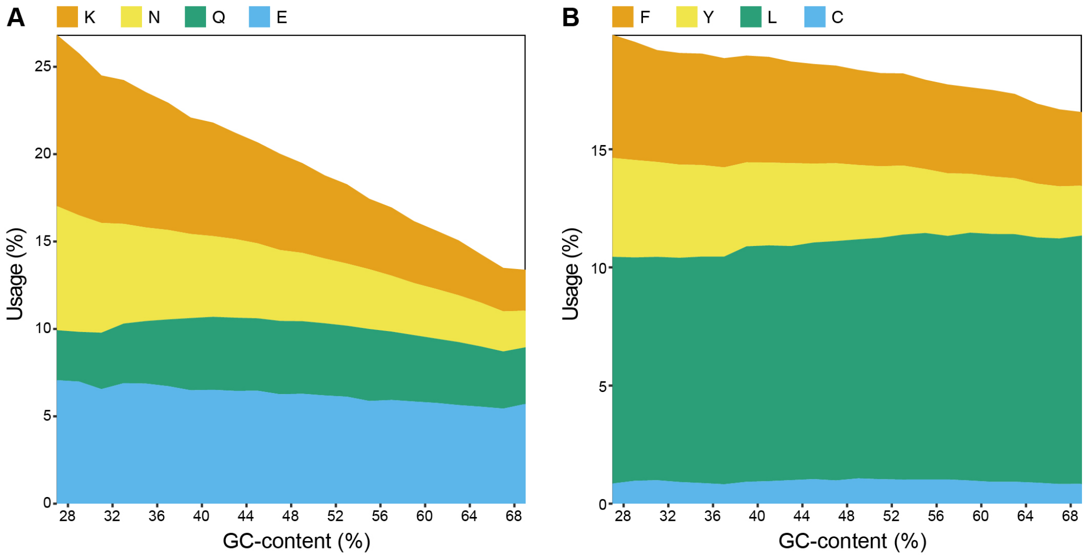


**Figure S12** Comparison between the usage of A-rich (A) and T-rich (B) amino acids.

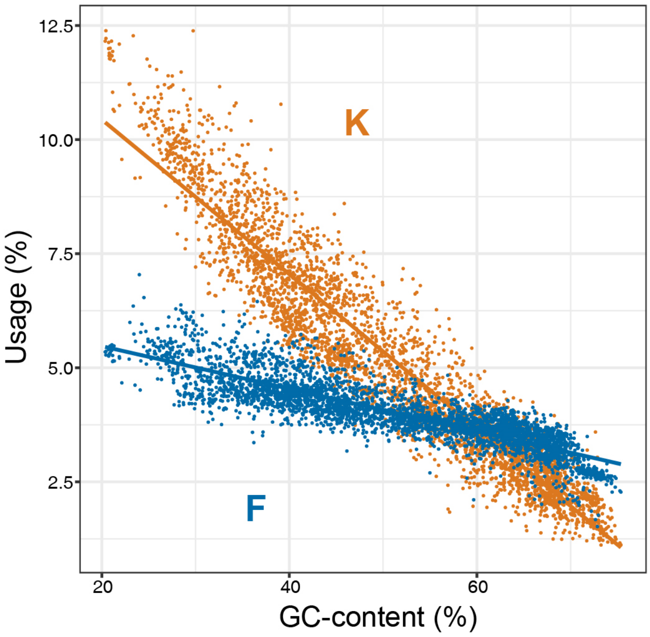


**Figure S13** Comparison between the usage of Lysine (K) and Phenylalanine (F) with different values of GC-content.

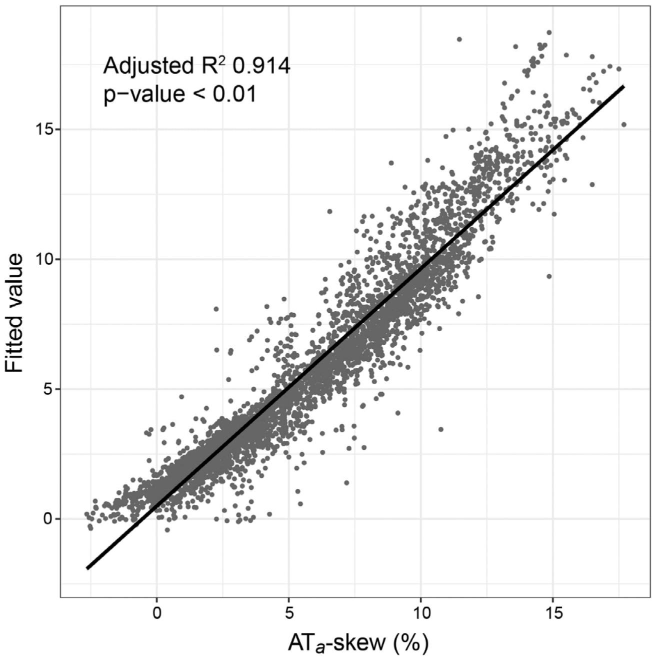


**Figure S14** Results of multiple regression analysis, where $AT_a$-skew was explained by the usage of Lysine (K) and Phenylalanine (F). Fitted values of $AT_a$-skew are shown in contrast to real values.

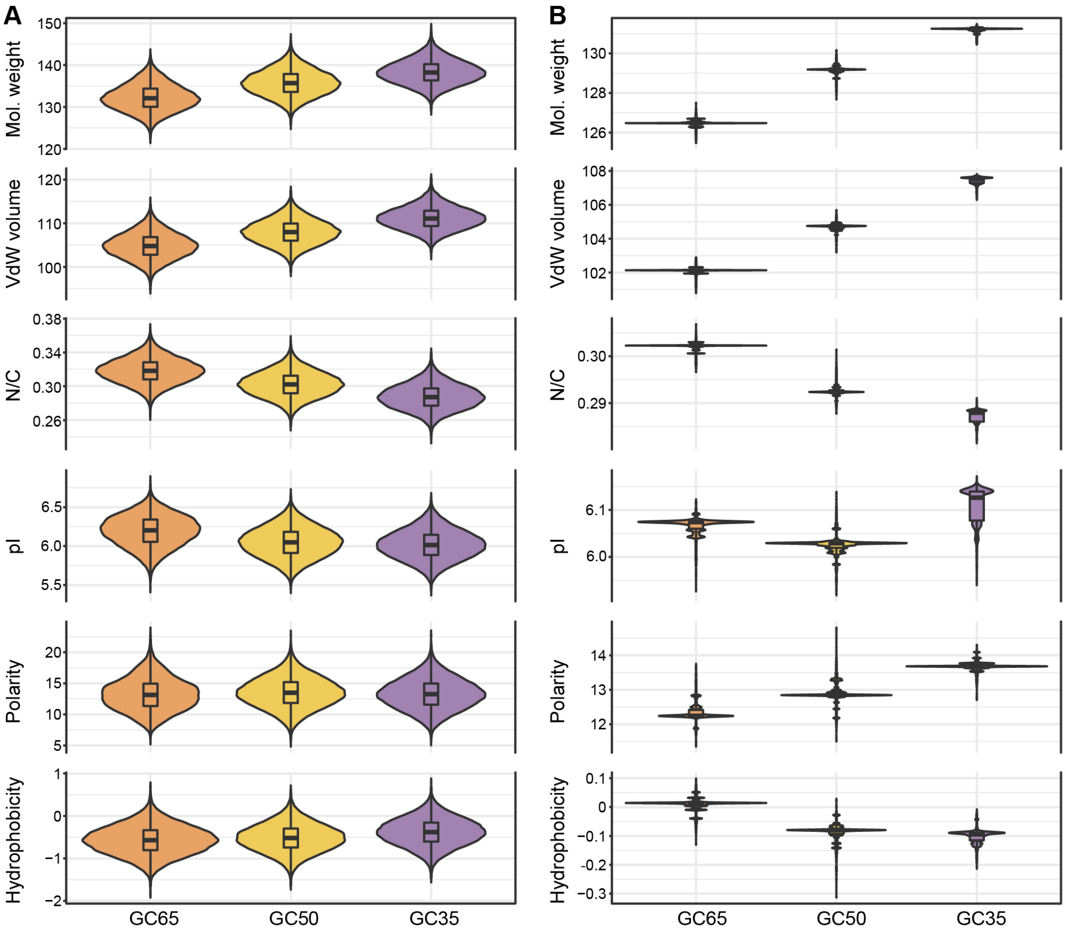


**Figure S15** Variations of average amino acids properties in established neutral (A) and natural (B) modes with different levels of GC-content.

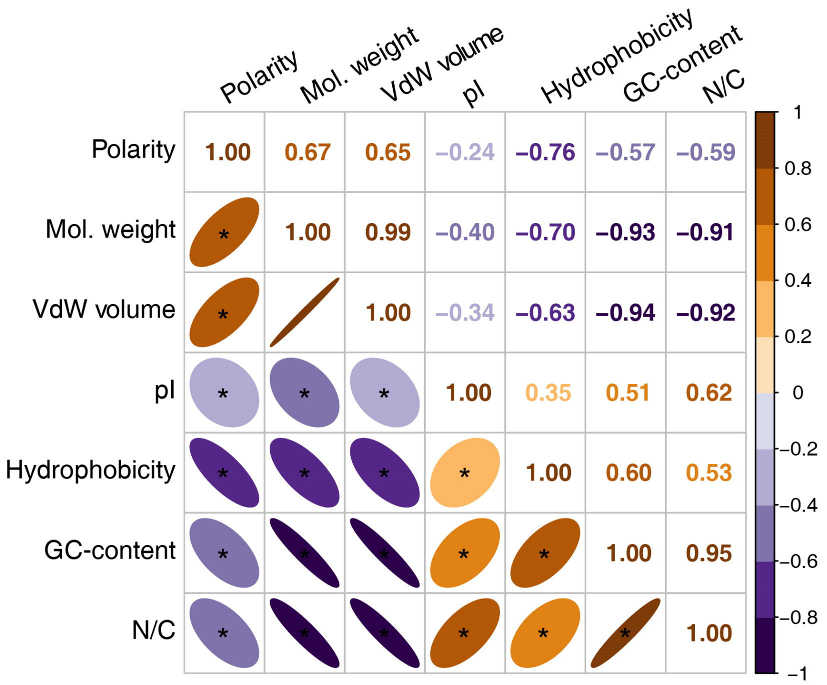


**Figure S16** Pairwise correlation coefficients between GC-content and average amino acids properties. Asterisks in the plot indicate adjusted p-value < 0.01.

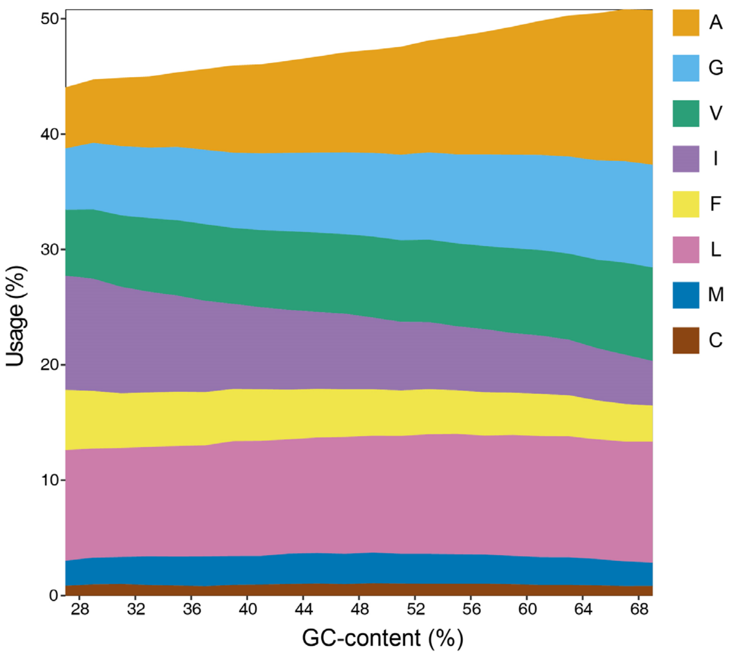


**Figure S17** Usages of the most hydrophobic amino acids ((i.e., I, L, V, F, C, M, A and G)) with different values of GC-content.
